# Multi-material biomaterial model of scaffold-defect integration at the wound margins

**DOI:** 10.64898/2026.03.10.710896

**Authors:** Alison C. Nunes, Grace Rubino, Huamin Gao, Megan Shamsi, Vasiliki Kolliopolous, Aleczandria Tiffany, Brendan A.C. Harley

## Abstract

Critical-sized craniomaxillofacial (CMF) defects affect the skull, face, and jaw, arising from conditions such as cleft palate, oncologic resections, and high energy impacts, and due to their large size and irregular geometry, cannot heal naturally by the body, thus requiring surgery. The field of biomedical research has long recognized the need to develop higher order biomaterial model systems for improved disease characterization and translational therapeutic/material progress. There is, however, difficulty in developing these workflows at the scale of conventional two-dimensional cell culture screening systems while simultaneously approaching a level of complexity necessary to consider translation to *in vivo* animal models. Here, we describe a three-dimensional (3D), *in vitro* model system to investigate the impact of stromal cell migration from one microenvironment to another at a medium-throughput scale. Importantly, we demonstrate the ability of this workflow to be utilized as a screening tool for collagen-based biomaterial motifs of interest in promoting craniomaxillofacial bone defect repair. Taken together we provide a strategy for interpreting cell-to-cell, cell-to-material, and material-to-material interactions across a multidimensional spatiotemporal scale.

## 1. Introduction

Critical-sized craniomaxillofacial (CMF) defects are particularly complex bone defects affecting the skull, face, and jaw that present a challenge for regenerative healing as they are typically large, and unique in shape. These can result from congenital, post-oncologic, or traumatic injuries [1–3]. Cleft palate is a highly prevalent example of a congenital CMF defect, occurring in 1 of every 1,600 babies born in the United and 26% of battle-field injuries in Iraq and Afghanistan involved penetrating CMF defects [1]. Current treatments rely on autografts (from the patient) or allografts (typically from a cadaver), which are limited by donor availability, immune rejection, and poor osteogenic potential. Alloplastic implants, another alternative, provide mechanical stabilization, but often do not degrade and require a secondary removal surgery [3–9]. Successful repair depends on rapid vascularization, driven by immune-mediated recruitment of endothelial cells to establish networks that support nutrient transport and implant integration [10]. Biomaterials pose a promising alternative to traditional treatment methods but must overcome two fundamental hurdles in the regenerating critical-sized CMF defects: (1) ensuring proper graft vascularization and (2) reducing the host inflammatory response, both of which are required to ensure successful implant integration. Therefore, a biomaterial implant that could drive processes involved in regeneration of these defects would be transformative for CMF repair.

Biomaterial implants for CMF repair offer an opportunity to optimize mechanics and biologic signaling required to promote tissue regeneration of these defects. The main advantage of biomaterials is that they are tailorable, allowing for the manipulation of multiple material properties to drive vascularization of the implant and bone growth. Current options of biomaterials are primarily space filling, with few options for true bone regeneration as well as difficulties in ensuring proper integration of the graft into the wound site due to inadequate vascularization and improper fitting of the implant to the wound margin due to geometric heterogeneity. Porous type I collagen scaffolds combined with glycosaminoglycans have been shown to successfully repair tendon and skin, as well as aid in bone repair with the addition of mineral-phase calcium and phosphate [11–19]. Benefits of this material include their unique, tunable pore architecture and orientation, their ability to retain growth factors, and most importantly the ease of incorporating additional materials during fabrication to manipulate specific implant properties [11, 12, 15, 20–23]. This combination of composition and chemistry promote human mesenchymal stem cells (hMSC) to deposit mineral *in vitro* and regenerate bone *in vivo* [24]. In larger scale defect model systems with mineralized collagen implants, we see inadequate vascular development and poor integration [25]. The most recent advances to address the gaps in healing potential of collagen scaffolds involve the development of composite materials that incorporate 3D-printed meshes such as polycaprolactone (PCL) and polylactic acid (PLA), allowing for improved mechanical stability and shape-fitting in heterogeneous wound margins as well as improved osteogenic potential [26–28]. However, the biologic complexity of the wound margin as well as heterogeneity in patients including diminished healing due to ageing or exposure to radiation in the case of oncologic resections, still presents problems for implant integration due to a lack of understanding of cellular behavior at the wound-implant margin, and a method for studying this complex interplay is critical in finally addressing this gap in knowledge.

Traditional models for studying patient heterogeneity have primarily been either two-dimensional (2D) cell culture platforms or animal models. 2D cell culture platforms allow for a narrow investigation of cellular behaviors and can be transformed to a high-throughput platform via the use of spheroids to answer specific cellular questions such as understanding attachment patterns due to various biomaterials, but do not capture the complexity of the native microenvironment or signaling between cells and the extracellular matrix (ECM) [29, 30]. Animal models allow for the use of a physiologically relevant environment but are limited in the ability to decouple processes due to the complex nature of the system. Therefore, the development of a well-controlled 3D microenvironment model that allows for increased complexity while maintaining the ability to identify and measure specific factors that may be influencing cell behavior is of critical importance in ensuring the translational potential of experimental findings. Current state-of-the-art 3D material model systems typically rely on some variation of separation between microenvironments like a microfluidic device. With this work we highlight the importance of direct contact of these microenvironments and establish a method to understand long-term feedback loops between these areas.

Here, we demonstrate an *in vitro* 3D composite biomaterial system that characterizes early processes of human mesenchymal stromal cell migration and differentiation into a porous scaffold from a precultured tissue mimetic hydrogel. As proof-of-concept development, we capture hMSC biologic responses to MC-GAG scaffolds of diameters ranging from 2mm-6mm grafted into this system for a six-day period after defect induction. Subsequently, we explored the osteogenic, immunomodulatory, and vascular hMSC bias resulting from varied glycosaminoglycan content within this mineralized collagen material at an extended timescale. These studies consider the importance of understanding early cellular activity into material grafts to promote regenerative healing for complete biomaterial integration post-implantation. We believe our injury workflow will serve as a powerful tool for screening new materials of interest at scale and provide an opportunity to directly exploit biologic questions associated with 3D multi-microenvironment systems and feedback loops.

## 2. Materials and Methods

### 2.1. Experimental Design

This study aimed to develop a methodology to understand hMSC migratory and osteogenic behavior patterns in an *in vitro* craniomaxillofacial defect model as a tool to screen biomaterial design motifs in medium throughput. Human mesenchymal stem cells were cultured in a gelatin-methacrylate (gelMA) hydrogel and a cylindrical defect was biopsy punched away. An acellular mineralized collagen scaffold was then implanted into the cylindrical defect site. Initial experiments sought to establish that we could quantify the migration of hMSC’s from the hydrogel to the scaffold graft after a one-day hydrogel culture period and subsequent seven-day graft period. DNA was isolated from the scaffolds to demonstrate migration into the graft and Ki-67 immunofluorescence staining was utilized to determine changes in cellular proliferation because of the graft and as a function of distance from the injury site. Next, an eight-day hydrogel culture period and subsequent six-day graft period was run to assess the role of defect/scaffold size on key regenerative indicators and migratory patterns via bulk model secreted factor assays and gene expression analysis in both the material graft and outer hydrogel shell. A secondary *in vitro* experiment was performed addressing the role of glycosaminoglycan content as a function of the same assay parameters for an extended 28-day period post defect/graft point and after a seven-day preliminary hydrogel culture period. Mineralized collagen scaffolds containing either heparin, chondroiton-4-sulfate (C4S), or chondroitin-6-sulfate (C6S) were grafted into 4mm cylindrical defects at the center of these hydrogels. Across this timeline, media was collected and analyzed for OPG and VEGF secretion quantification and RNA was similarly isolated to understand gene expression profiles.

### 2.2. Materials Fabrication

#### 2.2.1. Preparation of methacrylamide-functionalized gelatin (gelMA) and Characterization

Methacrylamide functionalized gelatin (GelMA) was synthesized utilizing previously described methods, 2g of type A porcine gelatin, 300 bloom (Sigma Aldrich) was dissolved in 20mL of sodium carbonate-bicarbonate buffer (Sigma Aldrich) of pH 9.4 at 50°C [31–34]. A dropwise addition of 80uL of methacrylic anhydride (Sigma Aldrich) was reacted with the previously made solution for 1 hour and 15 minutes. 80mL of deionized water at 50°C was added to stop the reaction. A seven-day dialysis period in 12-14kDa membranes took place with daily deionized water changes. The product was collected, frozen down for 24 hours and lyophilized for seven days. To confirm degree of functionalization (DOF) 1HNMR was utilized. *In vitro* work pooled multiple batches of GelMA together with a DOF range 0.48-0.59. Fabricated hydrogel mechanics were also assessed via an Instron 5943 mechanical testing apparatus (Instron). Hydrogels of 100uL volume were fabricated and tested under confined compression at 0.1mm/min to generate a stress strain curve from which we could obtain the Young’s modulus from the linear region of the stress-strain curve via an in-house MATLAB (Mathworks) code.

#### 2.2.2. Preparation of mineralized collagen-glycosaminoglycan scaffolds

Mineralized collagen-glycosaminoglycan scaffolds were fabricated by combining a collagen-mineral solution with the addition of either chondroitin-6-sulfate (C6S), chondroitin-4-sulfate (C4S), or Heparin (Hep) using previously mentioned protocols [24, 25, 35]. The following glycosaminoglycans (0.84 w/v%, C6S: Chondroitin sulfate sodium salt, Spectrum Chemicals, C4S: Chondroitin sulfate sodium salt, Sigma Aldrich, or Hep: Heparin sodium salt from porcine intestinal mucosa, Sigma Aldrich) were mixed with calcium salts (calcium hydroxide and calcium nitrate tetrahydrate, Sigma Aldrich), Type I bovine collagen (1.9 w/v% Collagen Matrix Inc.), and a solution of mineral buffer (0.1456 M phosphoric acid/0.037 M calcium hydroxide). Post homogenization, the resulting solution was added to square molds and lyophilized in a Genesis freeze-dryer (VirTis). A preset protocol was run to cool the suspension at a rate of 1°C/min from 20°C to -10°C. The resulting molds were left for an additional 2h at -10°C before being brought to 0°C and 0.2Torr to sublimate out the ice crystals. The resulting material is a solid collagen-GAG network with pores where the ice crystals had previously formed. Individual grafts were biopsy punched from the material using 2mm, 4mm, and 6mm biopsy punches (Integra LifeSciences).

#### 2.2.3. Collagen-glycosaminoglycan scaffold sterilization, crosslinking and hydration

Individual biopsy-punched scaffolds were placed in sterilization pouches and sterilized in an AN74i Anaprolene gas sterilizer (Andersen Sterilizers Inc.) for a 12-hour ethylene oxide cycle. Prior to grafting, sterile constructs were soaked in a 2-hour 100% sterile ethanol solution, washed and soaked in sterile 1x phosphate buffered saline (PBS), underwent an EDC-NHS crosslinking reaction in a 5:2:1 ratio of collagen:EDC:NHS for 1.5 hours, washed and soaked in PBS, and soaked in basal growth media for 48 hours. This hydration and crosslinking follow previously described methods [24, 35].

### 2.3. Cell Culture

#### 2.3.1. Cell preparation and materials preparation

Human bone-marrow-derived mesenchymal stem cells of a 26-year-old female (Essent Biologics) were expanded from passage 3 to passage 4 utilizing RoosterNourish expansion media (RoosterBio) for 3 days of flask culture. On the third day, media was replaced with low glucose DMEM and glutamine, 10% mesenchymal stem cell fetal bovine serum (Gemini), and 1% antibiotic-antimycotic (Gibco) for 2 days of culture. Flasks were maintained in aseptic incubation conditions at 37°C and 5% CO_2_. On day 5 of culture, cells were lifted from T175 flasks using TrypLE Express Enzyme (Thermo Fisher Scientific) for 5 min under 37°C incubation and 5% CO_2_. Culture media was utilized to neutralize the TrypLE and the cells were pelleted via centrifugation at 200g for 10 minutes. The TrypLE+media supernatant was removed via aspiration and cells were resuspended in media, counted (Countess Automated Cell Counter, Invitrogen) and distributed to individual stocks for hydrogel resuspension.

#### 2.3.2. Fabrication of cell embedded gelMA hydrogels

A methacrylated gelatin suspension (5wt% gelMA in 1x PBS with 5wt% Photoinitiator) was utilized to resuspend a cell pellet of hMSC to generate a precursor suspension at 1x10^6^cells/mL. The precursor was pipetted into a 48-well plate and photopolymerized for 1 minute with a UV lamp (Digital Light Lab) to generate 2mm height cylindrical gels confined to the well plate. Growth media, in 500uL increments, was added to each hydrogel and monitored via incubation under 5% CO_2_ at 37°C prior to biopsy punching and graft implantation.

#### 2.3.3 Hydrogel biopsy punching and scaffold implantation

Following the 6-day preculture period, biopsy punches (Integra) of diameters 2, 4 and 6mm were utilized to generate cylindrical defects within the precultured hydrogels on day 7 of culture. Cylindrical defects were collected for preliminary analysis and mineralized collagen scaffold grafts were placed into the open sites.

### 2.4. Viability

Cellular DNA was isolated using a Zymo DNA isolation kit (Zymo Research Corporation) per manufacturers protocol. A concentration reading was gathered using a NanoDrop Lite Spectrophotometer (Thermo Fisher Scientific) and a representative cell number measurement was quantified from a standard linear curve generated on day 0.

### 2.5. Isolation of RNA and NanoString nCounter Analysis Pipeline

RNA was separately isolated from each model component (punch, scaffold, and outer hydrogel) across the 14 and 28-day time courses utilizing a Zymo RNA Isolation Kit per manufacturer’s instructions (Zymo Research). Upon elution, RNA concentration was measured via NanoDrop Lite Spectrophotometer (Thermo Fisher Scientific). A custom NanoString panel of mRNA probes was utilized to quantify the transcript expression level using a NanoString nCounter System at the Tumor Engineering and Phenotyping Shared Resource at the Cancer Center at Illinois. RNA was loaded into the cartages and run according to the manufacturers protocol. Available nSolver analysis software and Excel were utilized for data processing and normalization.

### 2.6 Cell secreted factors collection and analysis

Media was collected from each model and refreshed every day across the entire 14-day and 28-day experiments. Enzyme linked immunosorbent assays (R&D Systems) were used to quantify the amount of VEGF and OPG released across the whole model system at timepoints of interest. Samples were processed according to manufacturer’s protocol and absorbances were read on an Infinite Tecan system (Tecan). Each analyte had background absorbance subtracted out and was converted to a concentration utilizing a known concentration curve. Values are reported as cumulative concentration.

### 2.7. Statistics

Statistics were performed utilizing GraphPadPrism (USA) and Microscoft Excel (USA) software. For all analyses, significance was set to p<0.05. QQ plots were generated to assess if the residuals are sampled from a Gaussian distribution. If applicable, comparisons between multiple groups were performed utilizing a two-way ANOVA. A Tukey multiple comparisons test was used to compare significance (p<0.05) between two groups of interest. Technical and biologic replicates were included in all quantitative analysis.

## 3. Results

### 3.1 hMSC’s migrate from surrounding hydrogel into scaffold within seven days

We first performed preliminary experiments to determine whether we could quantify hMSC migration from the hydrogel into the scaffold graft over the course of 7-days. DNA quantification showed an average of approximately 4500 cells migrated into the grafts over the 7-day period (Fig. 1).

**Fig. 1.**
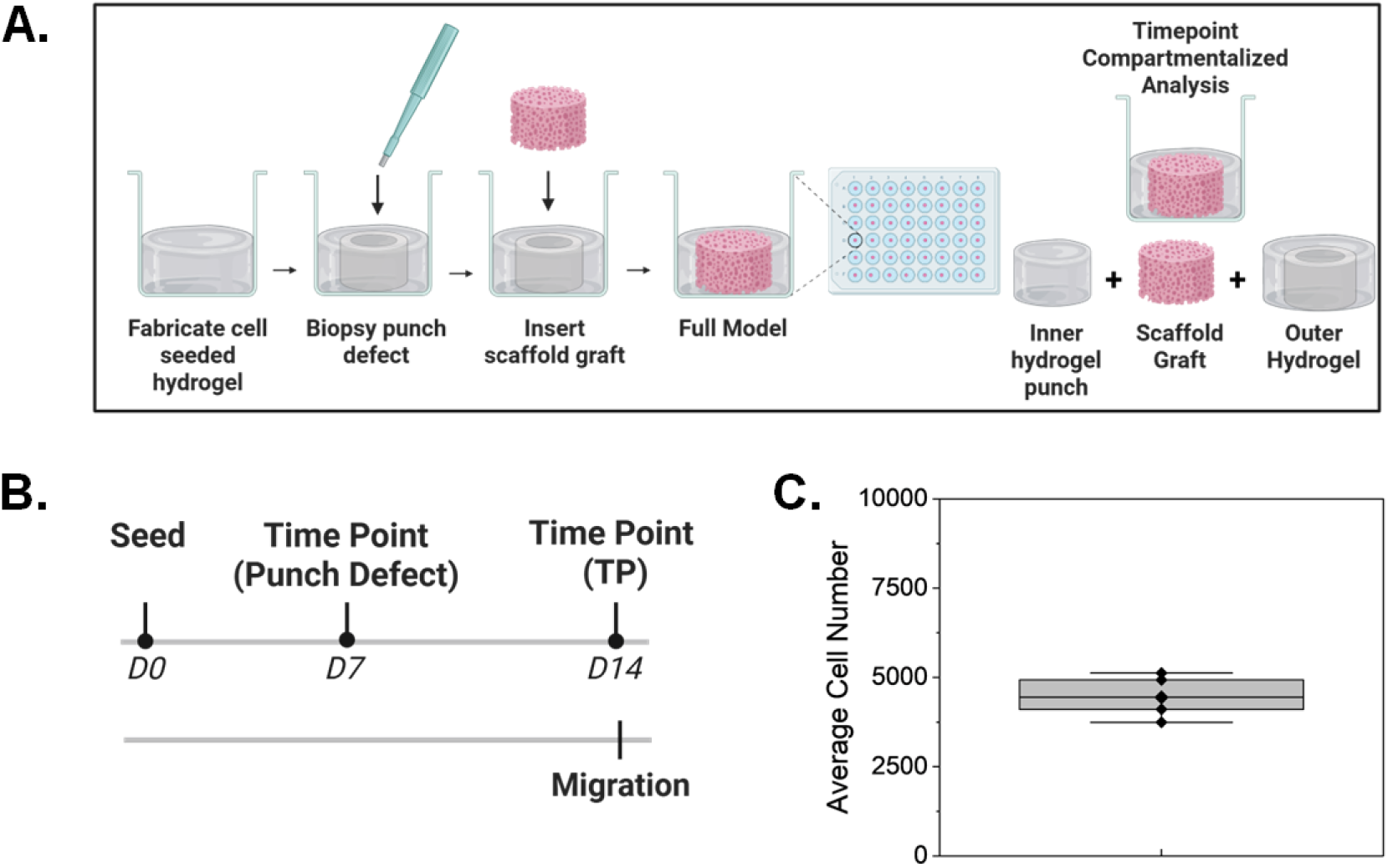
**A)** Conceptual overview of the wound model. **B)** Experimental plan for a 7 day hMSC migration experiment into implanted scaffold. **C)** Mean cell number found withgin implanted 4mm dia. scaffold after 7 days in culture.

### 3.2 Observed effects of scaffold glycosaminoglycan content on MSC secretome within the wound model

We then aimed to assess the ability of our platform to identify differences in biomaterial constructs via the incorporation of different glycosaminoglycans, which have previously been shown to modulate hMSC gene expression when directly seeded onto GAG scaffolds (Fig. 2). Though not significant, an average increase in VEGF and OPG release from hMSC over time was observed across all model systems. At day 28, we observed significantly higher levels of secreted VEGF from model systems containing scaffolds in comparison to the full hydrogel model. hMSC secreted OPG was significantly higher from both the bulk hydrogel model and the model containing heparin compared to models with C6S and C4S scaffold grafts. A jump in OPG release across all model systems is observed from day 14 to day 21 and again from day 21 to 28 where a significant increase in hMSC secretion is observed.

**Fig. 2.**
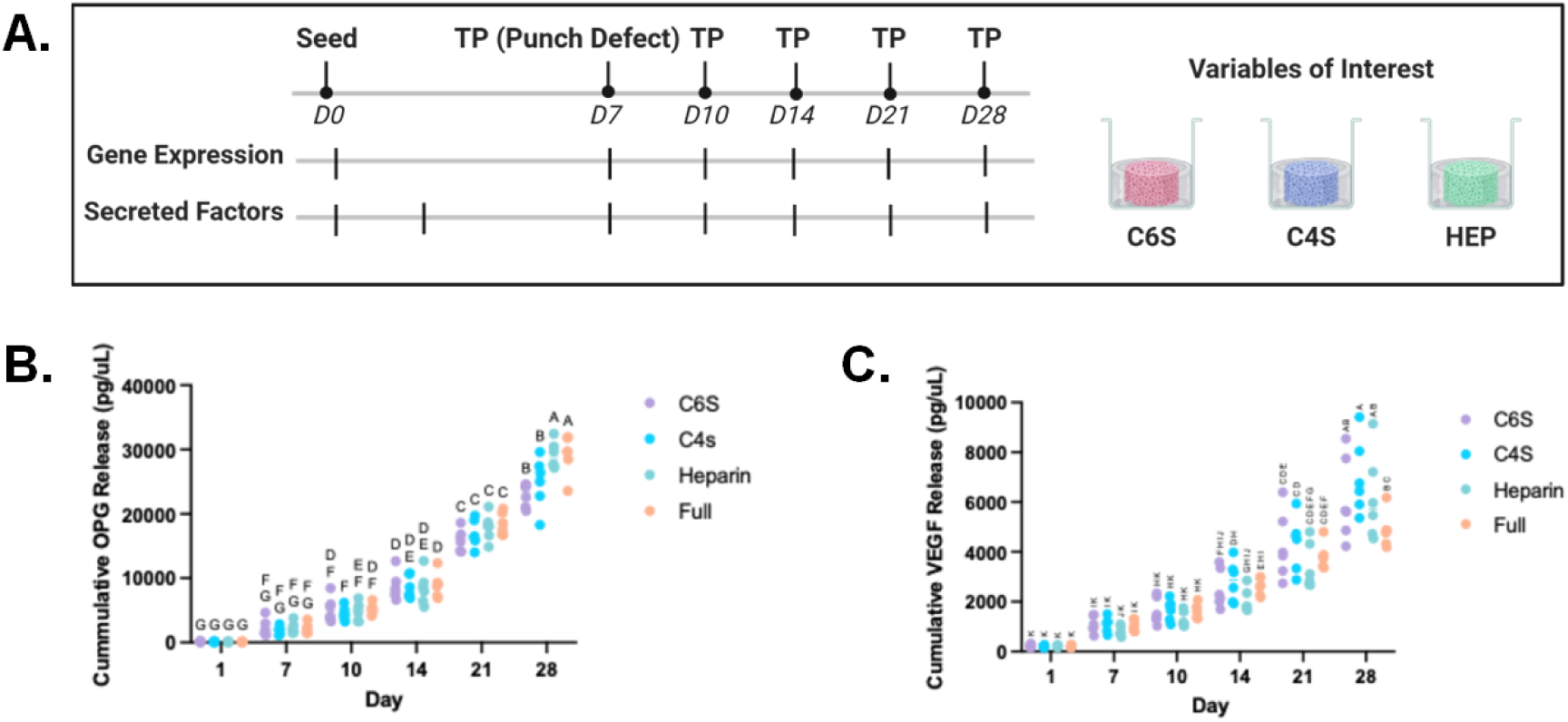
**A)** Experimental plan to assess hMSC recruitment and activity in mineralized collagen scaffolds as a function of incorporated GAG content. **B)** Endogenous production of osteoprotegerin (OPG) by recruited hMSCs within the scaffold as a function of scaffold GAG content. **C)** Endogenous production of VEGF by recruited hMSCs within the scaffold as a function of scaffold GAG content. **Statistics:** groups sharing the same letter do no differ significantly (p < 0.05).

### 3.2 Scaffold glycosaminoglycan content influences hMSC transcriptomic signature within the wound model

Gene expression analysis was performed to assess fold change differences in hMSC transcriptional levels at day 10 and day 28 in the scaffolds compared to day 0 controls (Table 1, Fig. 3). Three days post graft placement, IDO1 is significantly upregulated in C6S scaffold constructs, OPN is upregulated in C4S scaffold constructs, and IHH is upregulated in heparin scaffolds (Fig. 3). Notably no genes were significantly downregulated. By day 28, IL1RN was upregulated in C4S scaffolds and SOX9 was upregulated in heparin scaffolds (p < 0.05) (Fig. 3). HGF was significantly downregulated across all scaffold variants by day 28 and ALPL was significantly downregulated in hMSC migrated to both C4S and heparin containing scaffolds (Fig. 3).

**Table 1.** Complete list of all regenerative genes evaluated within the scaffold implant at days 10, 28 post implantation. Red/blue corresponds to relatively up- or down-regulated versus hMSCs at day 0. *: p < 0.05. **: p < 0.01.

| Gene | Category | D10 | D28 |
| --- | --- | --- | --- |
| *ANGPT1* | Angiogenic | * |  |
| *VEGFA* | Angiogenic | * |  |
| *ALPL* | Osteogenic | * |  |
| *BGLAP* | Osteogenic | * |  |
| *BMP2* | Osteogenic |  | * |
| *BMP7* | Osteogenic |  | *** |
| *COL1A2* | Osteogenic |  | * |
| *SEMA3A* | Osteogenic |  | * |
| *SP7* | Osteogenic |  | *** |
| *TNFRSF11A* | Osteogenic | ** | ** |
| *TNFRSF11B* | Osteogenic | * |  |
| *WNT16* | Osteogenic |  | ** |
| *CCL7* | Immunomodulatory |  |  |
| *IDO1* | Immunomodulatory |  | ** |
| *IL10* | Immunomodulatory |  | * |
| *IL1RN* | Immunomodulatory |  | ** |
| *PTGS2* | Immunomodulatory | * |  |
| *TNFSF11* | Immunomodulatory | * | ** |

**Fig. 3.**
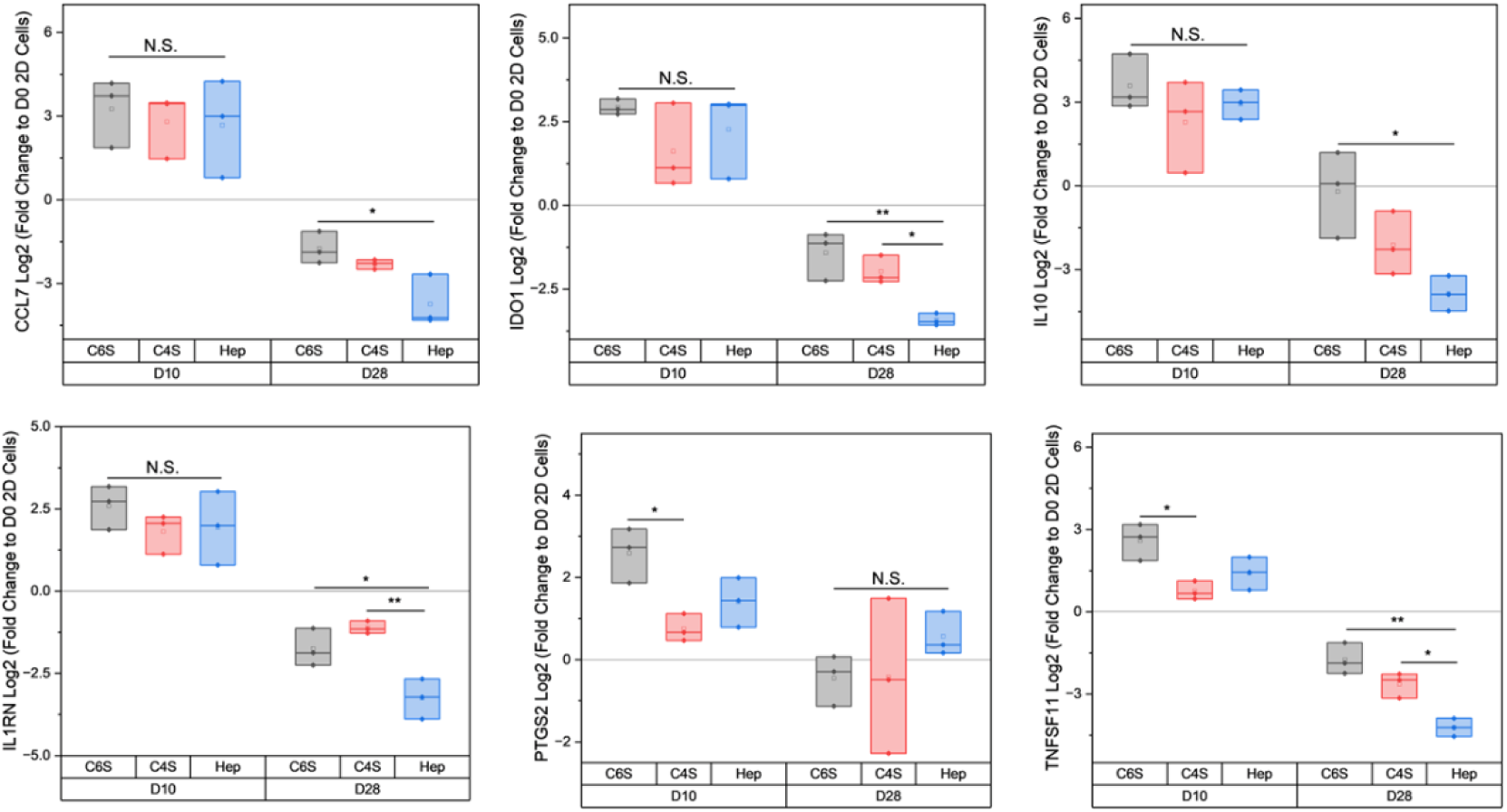
Differentially regulated immunomodulatory genes for hMSCs recruited into implanted mineralized collagen scaffolds from the surrounding hydrogel defect at day 10 or 28 post scaffold implant. *: p < 0.05. **: P < 0.01.

We also examined changes in gene expression patterns in the hydrogel wound as a result of the implantation of scaffolds into the defect, reasoning the secretome of cells in the scaffolds could alter the activity of hMSCs that remained in the surrounding hydrogel wound margin (Table 2, Fig. 4). At day 10 no genes were upregulated in hydrogels of the C4S, HEP, or full models. Human mesenchymal stem cells seeded in C6S containing models saw significantly upregulated levels of SOX9 and significantly downregulated levels of HGF at day 10. Scaffold containing model systems outer hydrogels saw no significant downregulation of gene levels at day 28, while the bulk hydrogel control saw a significant decrease in HGF levels. IGF2 was significantly upregulated in outer hydrogels of models with both C6S and heparin containing scaffolds at day 28. Additionally, VEGFA was significantly upregulated in C6S outer hydrogels and FGFR2 was significantly upregulated in outer hydrogels of models containing heparin scaffolds.

**Fig. 4.**
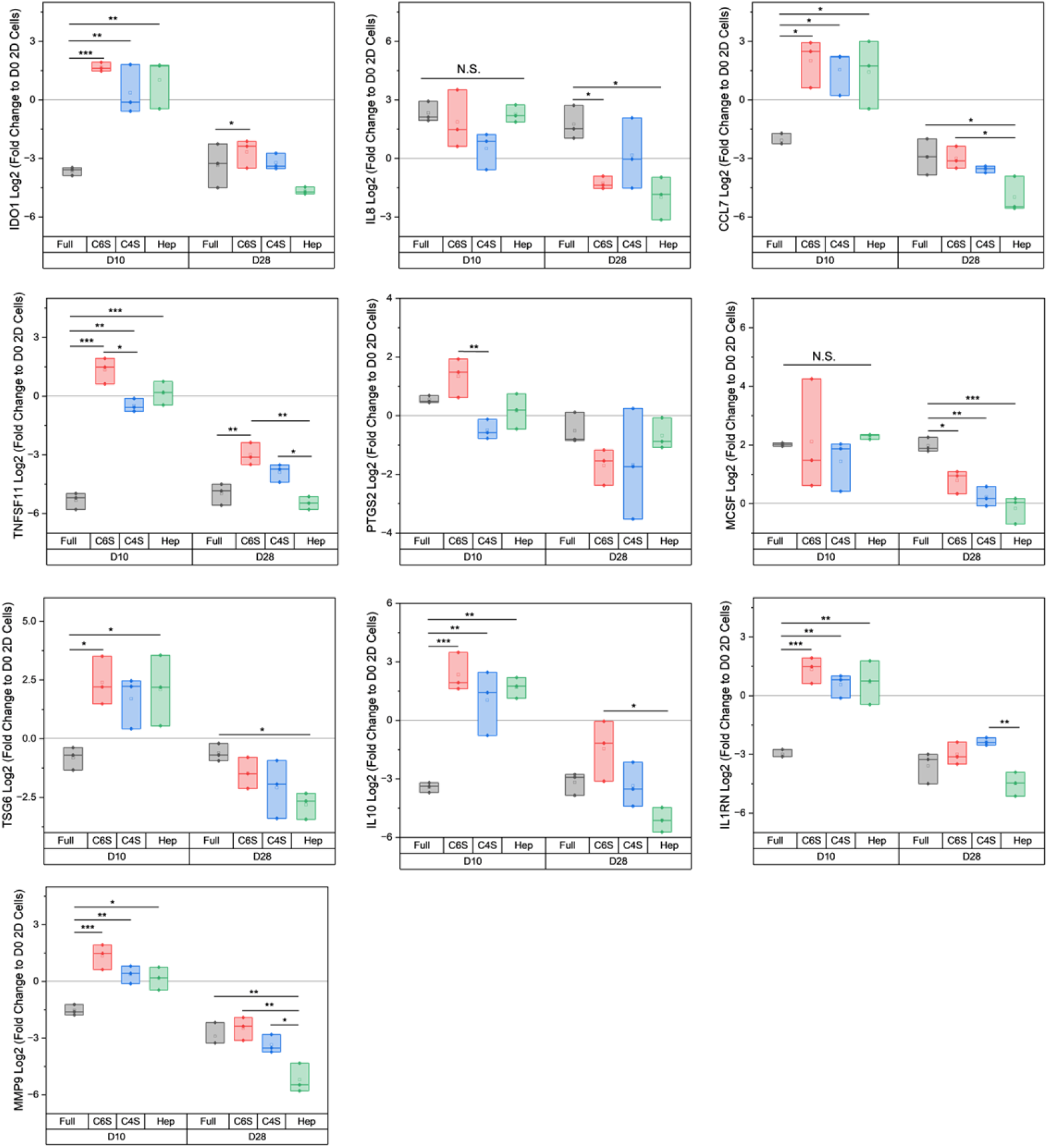
Differentially regulated genes for hMSCs that remained in the outer hydrogel wound model as a function of implanted mineralized collagen scaffold glycosaminoglycan content at day 10 or 28 post scaffold implant. Comparisons normalized to day 0 hMSCs with results for each scaffold implant composition also compared to non-injured hydrogels (Full). *: p < 0.05. **: p < 0.01. **: p < 0.001.

**Table 2.** Complete list of all regenerative genes evaluated within the surrounding hydrogel margins at days 10, 28 post implantation. Red/blue corresponds to relatively up- or down-regulated versus hMSCs at day 0. *: p < 0.05. **: p < 0.01. ***: p < 0.001.

| Gene | Category | D10 | D28 |
| --- | --- | --- | --- |
| *ANGPT1* | Angiogenic | *** | *** |
| *GAL9* | Angiogenic | *** |  |
| *HGF* | Angiogenic |  | ** |
| *VEGFA* | Angiogenic | *** |  |
| *VEGFB* | Angiogenic |  | * |
| *ALPL* | Osteogenic | *** | ** |
| *BGLAP* | Osteogenic | *** |  |
| *BMP2* | Osteogenic | *** | ** |
| *BMP7* | Osteogenic | *** | *** |
| *COL1A2* | Osteogenic |  | ** |
| *IHH* | Osteogenic | *** | * |
| *OPN* | Osteogenic | *** |  |
| *RUNX2* | Osteogenic | ** |  |
| *SEMA3A* | Osteogenic | ** | ** |
| *SMAD5* | Osteogenic |  | ** |
| *SP7* | Osteogenic | *** | * |
| *TNFRSF11A* | Osteogenic | *** | ** |
| *TNFRSF11B* | Osteogenic | *** |  |
| *WNT16* | Osteogenic | *** | *** |
| *SOX9* | Chondrogenic |  | * |
| *CCL7* | Immunomodulatory | * | * |
| *IDO1* | Immunomodulatory | *** | *** |
| *IL8* | Immunomodulatory |  | * |
| *IL10* | Immunomodulatory |  | * |
| *IL1RN* | Immunomodulatory | *** | ** |
| *MCSF* | Immunomodulatory |  | *** |
| *MMP9* | Immunomodulatory | *** | ** |
| *PTGS2* | Immunomodulatory | ** |  |
| *TNFSF11* | Immunomodulatory | *** | ** |
| *TSG6* | Immunomodulatory | * | * |

## 4. Discussion

The multivariable complexities associated with biomaterial modeling in 3D increase the difficulty of designing long term high-throughput studies. Most high order model systems rely on microfluidic devices, particularly those incorporating two or more microenvironments of interest. These devices rely on channels, often of non-biologic material, with a perfusion flow that integrates the two environments together [36–41]. There is a significant opportunity to improve within the 3D biomaterial modeling space, to design systems that are better able to capture more physiologically relevant cellular level responses in higher throughput, ultimately helping to accelerate the pharmaceutical and biotechnology development pipeline.

In this study, we present a unique model system that aims to provide a methodology to enhance our understanding of cell kinetics toward a biomaterial or microenvironment of interest. Different from other 3D assays and models, this system incorporates a graft like mechanism providing a flexible opportunity to create an initial microenvironment of interest and understand time altered cellular responses to a new material environment. Here we demonstrate an array of design strategies for building a 3D biomaterial graft model, specifically accounting for the effect of graft size on hMSC migration, secretion, and differentiation capacity. Once established, we secondarily probed the effect of glycosaminoglycan content on hMSC migration, differentiation, transcriptomic and proteomic responses. Overall, we observe subtle influence of graft surface area to volume ratio on functional outputs, both transcriptomic and proteomic. Secondarily we define consistent trends in assay readouts associated with varied glycosaminoglycan content in scaffold graft constructs to previous studies. Lastly, we explore new questions aligned with attempts to understand feedback loops in a spatial temporal context between separate but in contact microenvironments that are evolving in time.

As a first pass study, we aimed to determine the effect of defect/graft size on hMSC migration kinetics, viability, and transcriptional and proteolytic expression. In particular, we probed three defect/graft sizes of interest: 2mm, 4mm, and 6mm diameter. Notably, defect/graft size impacts cell density and metabolic activity within the scaffold post grafting, but does not impact outer hydrogel hMSC density or metabolic activity. Significant differences in the release of soluble osteogenic factors (VEGF and OPG) between the 6mm defect/graft model system and all other model systems were observed. We attribute this difference to be due to the largest defect removal and infer that too many hMSC were lost via the creation of the defect to observe adequate migration over the experimental timescale. When normalized to a day 0 transcriptional level, we observe a significant downregulation of HGF at day 14 across all model scaffold graft variants and no significance with respect to these gene in the bulk hydrogel control, indicating that the scaffold is driving transcriptional changes over time different from the hydrogel. Additionally, common trends in genes up and downregulated across the size variants gives confidence that surface area to volume ratio of defect does not highly impact transcriptional level changes. Follow up work utilizes the 4mm defect design which targets the greatest user feasibility, maintains lowest possible material usage, and consistent cellular level functional responses and outputs. While this work focuses on downstream hMSC activity after a mock craniofacial graft event, there remains an opportunity to apply this model system to any biologic or biomaterial context of interest within the constraints of the investigated scales.

Glycosaminoglycan is a majority component of bone extracellular matrix. Previously, mineralized collagen scaffolds with C6S, C4S, and heparin have been investigated from an hMSC osteogenic differentiation lens, a hUVEC angiogenic potential, and from a macrophage polarization perspective [42–46]. Observably, C6S containing scaffolds promote hMSCs to deposit calcium and phosphorous compared to C4S and heparin containing scaffolds in addition to promoting significantly greater network length formation in hUVEC. We observe similar responses in this work to previous work, showing that scaffold grafts in isolation cultured with hMSC, both in the presence of a licensed immunomodulatory stimuli and a basal control stimuli, secreted higher levels of OPG. With the defect model, systems containing heparin scaffolds saw a significantly higher level of OPG secreted at day 28 compared to other scaffold variants (Fig. fkdsl;). Additionally we observe a significant difference in VEGF release at day 28 from all model systems containing scaffolds compared to the bulk hydrogel control, indicating that the scaffolds are indeed driving a difference in response to the singular microenvironment with a particular vascular bias. Notably, we further probe the relationship of glycosaminoglycan content on hMSCs in a gelatin based neighboring microenvironment. In some cases, C6S and C4S outer hydrogels, we see downregulation of HGF which is also observed in their scaffold microenvironment. A similar downregulation is observed in the bulk hydrogel control. Overall, significant upregulation in osteogenic genes within the hMSC migrated to heparin containing scaffolds compared to immunomodulatory genes in hMSC migrated to C4S containing scaffolds reaffirms previously obtained results in singular scaffold studies. Interestingly we observe a difference in the genes that are upregulated in the scaffold constructs compared to their outer hydrogel suggesting an influence of the resulting bias affected migrated hMSC embedded in within the scaffolds.

## 5. Conclusions

The goal of this work was to establish a framework for construction and evaluation of a 3D biomaterial graft model *in vitro*. Additionally, we explored the use of our migration model assay to contextualize scaffold variant motifs of interest to improve craniomaxillofacial defect repair. Observably, we found that consistent trends in hMSC transcriptional and proteolytic outputs were maintained across graft variants. Migration and metabolic levels varied due to post graft cell numbers. We report the addition of heparin to scaffold matrix promotes late stage hMSC metabolic activity, proliferation, and OPG release. Scaffolds containing C4S showed potential for inducing immunomodulatory effects from hMSC while C6S containing models promoted osteogenic activity of hMSCs. Taken together, we observe consistent trends from previous work investigating scaffold variants in isolation without the hydrogel shell giving rise to further validate this model system. Future directions include creating a co-cultured vascularized hydrogel to probe multicellular responses and the post graft angiogenic timeline into implants *in vitro.* Furthermore, we propose the investigation of new materials and biologic contexts where migration and component microenvironments sit at the height of scientific inquiry: tumor metastasis, drug delivery, and immune responses to highlight a few.

## Acknowledgements

The authors would like to acknowledge the following institutes for access to their facilities and services: the School of Chemical Sciences Microanalysis Laboratory, the Carl R. Woese Institute for Genomic Biology, the Tumor Engineering and Phenotyping Shared Resource (TEP) at the Cancer Center at Illinois, and the Beckman Institute for Advanced Science and Technology, located at the University of Illinois. This manuscript is the result of funding in whole or in part by the National Institutes of Health (NIH). It is subject to the NIH Public Access Policy. Through acceptance of this federal funding, NIH has been given a right to make this manuscript publicly available in PubMed Central upon the Official Date of Publication, as defined by NIH. Research reported in this publication was supported by the National Institute of Dental and Craniofacial Research of the National Institutes of Health under Award Number R01 DE030491 (BACH) as well as National Institute of Arthritis and Musculoskeletal and Skin Diseases under Award Number R01 AR077858 (BACH). We are also grateful for funds provided by the NSF Graduate Research Fellowship (DGE-174604 to VK; DGE-1144245 to AST) and the Chemistry-Biology Interface Research Training Program at the University of Illinois (T32 GM070421, VK). Additional support was provided by the Carl R. Woese Institute for Genomic Biology and the Chemical and Biomolecular Engineering Dept. at the University of Illinois at Urbana-Champaign. The interpretations and conclusions presented are those of the authors and are not necessarily endorsed by the National Institutes of Health or the National Science Foundation.

## Contributions (CRediT: Contributor Roles Taxonomy [47, 48])

**A.C. Nunes:** Conceptualization, Data curation, Formal Analysis, Visualization, Investigation, Methodology, Writing – original draft, Writing – review & editing. **G. Rubino:** Investigation, Methodology, Formal Analysis. **H. Gao:** Investigation. **M. Shamsi:** Investigation. **V. Kolliopoulos:** Conceptualization, Investigation, Methodology, Writing – review & editing. **A. Tiffany:** Conceptualization, Investigation, Methodology, Writing – review & editing. **B.A.C. Harley:** Conceptualization, Resources, Project administration, Funding acquisition, Supervision, Writing – review & editing.

